# Expression of type I interferon-associated genes at point of antiretroviral therapy interruption predicts time to HIV virological rebound

**DOI:** 10.1101/2020.11.24.395574

**Authors:** Panagiota Zacharapoulou, Emanuele Marchi, Ane Ogbe, Nicola Robinson, Helen Brown, Mathew Jones, Lucia Parolini, Matthew Pace, Nicholas Grayson, Pontiano Kaleebu, Helen Rees, Sarah Fidler, Philip Goulder, Paul Klenerman, John Frater

## Abstract

Although certain individuals with HIV infection can stop antiretroviral therapy (ART) without evidence of viral load rebound, the mechanisms under-pinning ‘post-treatment control’ remain unclear. Twelve individuals who had received 12 months of ART from primary HIV infection and then undertook a TI were sampled at the time of stopping therapy. Using RNA-Seq we explored gene expression in CD4 T cells to look for evidence of a mechanism that might underpin virological rebound and lead to discovery of an associated biomarker. Using independent analysis tools, genes associated with the type I interferon response were strongly associated with a delayed time to viral rebound following TI. These are the first data we are aware of that link transcriptomic signatures associated with innate immunity with control following TI. While these results need to be confirmed in larger trials, they could help define a strategy for new therapies and identify new biomarkers for remission.

## Introduction

Unless suppressed by antiretroviral therapy (ART), HIV infection leads to persistent viremia, progressive loss of CD4+ T cells and eventually AIDS. However ART is not a cure for HIV infection, requires lifelong adherence, and has been associated with side-effects, drug resistance and stigma(Peluso et al., 2019). Inherent to testing novel HIV cure strategies(Dash et al., 2019; Davenport et al., 2019; Mendoza et al., 2018) is analytical treatment interruption (TI) in which ART is stopped and any delay or prevention in viral rebound is examined. However there is still much controversy around TIs, and how to best implement and interpret them(Julg et al., 2019). Identifying biomarkers which predict outcomes following TI would provide enormous value to both understanding mechanisms of virological remission and identifying new drug candidates.

A rare proportion of ART-treated individuals are able to stop therapy and maintain undetectable viremia for months and, in some cases, years(Etemad et al., 2019; Martin et al., 2017; Sáez-Cirión et al., 2013). Multiple studies have been attempted to elucidate the biological mechanisms underlying these outcomes, and numerous clinical factors and molecular biomarkers which correlate with time to rebound have been proposed (Conway et al., 2019; Hurst et al., 2015; Krebs & Ananworanich, 2016; Martin et al., 2017; Sharaf et al., 2018; Stöhr et al., 2013; Williams et al., 2014). However, the host factors that affect the responsiveness of T cells have not been thoroughly characterised (Hyrcza et al., 2007), especially at a transcriptome level, possibly due to small cohort sizes and the availability of samples at key timepoints. Additionally, many gene expression studies have employed microarray technology(Lee et al., 2019; L.-L. Zhang et al., 2017) which may be associated with limitations compared with more recent RNA-Seq approaches(Zhao et al., 2014).

SPARTAC (Short Pulse Antiretroviral Treatment at HIV-1 Seroconversion)(Fidler et al., 2013) was one of the largest randomized clinical trials to study different durations of ART administered to participants with Primary HIV Infection (PHI) and the impact on disease progression. Previous analyses revealed that although the majority of participants experienced HIV viral rebound shortly after stopping ART (i.e <4 weeks post TI), some maintained undetectable viraemia to the end of the study (>500 days after TI). We studied participants who received 12 months of ART started in primary HIV infection (PHI), and analysed CD4+ T cell mRNA at the time of TI. Our aim was to identify genes or gene-sets expressed by CD4+ T cells that might associate with longer periods of remission after TI. Our results could be integrated into personalised diagnostic algorithms, to inform therapeutic strategies towards HIV cure and to better understand the mechanism of HIV latency and remission.

## Results

### Clustering of participants based on clinical response

We identified 18 SPARTAC participants who had received 12 months of ART commenced during PHI, and who had viable samples of PBMCs taken at the point of TI. Clustering analysis of RNA expression revealed distinct groups based on ethnicity and gender (See Table 1 and Supplementary Figure S1), which might have contributed to the clinical outcomes. Therefore, to avoid potential confounding effects, we excluded six individuals resulting in a study group comprising twelve women, all from South Africa. ‘Days to viral load rebound after TI’ was used to define clinical phenotypes and structure the comparative analysis (Table 1).

**Table 1:** Participant Demographics

|  |  | Original Cohort |  | Final Cohort |
| --- | --- | --- | --- | --- |
| Sex |  | Female<br>(n=14) | Male<br>(n=4) | Female<br>(n=12; 100%) |
| Rebound | Early (<100 days to rebound) | 5<br>(35.70%) | 2 (50%) | 4 (33.3%) |
|  | Intermediate<br>(>100 & <500days to rebound) | 4<br>(28.50%) | 2 (50%) | 3 (25%) |
|  | Late (> 500 days to rebound) | 5<br>(35.70%) | 0 | 5 (41.6%) |
| Country | Europe | 1 (7.10%) | 4 (100%) | 0 |
|  | Africa | 13<br>(92.80%) | 0% | 12 (100%) |

Based on observations in SPARTAC and other cohorts(Colby et al., 2018, 2020; Etemad et al., 2019; Stöhr et al., 2013) we defined ‘early’ rebounders following TI (<100 days) as representing normal progressors, ‘intermediates’ (100-500 days) as representing a phenotype reflecting transient remission, and ‘late’ rebounders (>500 days) to be more characteristic of elite or ‘sustained’ controllers. As these cut-offs (i.e 100 and 500 days) have not been previously ratified - and there might be some overlap between them - we tested their validity by dividing the dataset based on these phenotypes in two different ways. First, we grouped the ‘intermediates’ and ‘lates’ together as Post-Treatment Controllers (PTC) (i.e all rebound >100 days) and compared these with the Progressors (<100 days). Next, we grouped the ‘early’ and ‘intermediate’ groups (i.e all rebound <500 days) into a group of non-sustained controllers (non-SC) and compared these with the sustained controllers (SC; >500 days). (Table S1). For this dataset, grouping the “intermediates” with either “early” or “late” rebounders helps identify genes associated with the more extreme phenotypes (i.e early or late rebounders) and improves the sample sizes for comparison, but at the potential risk of increasing variability and noise.

### Differential Gene Expression and Gene Set Enrichment Analysis shows a strong association of type I interferon pathways with sustained control of viremia

Differential Gene Expression (DGE) analysis was performed to identify genes that were differentially expressed when comparing clinical groups. Genes with a reported adjusted p-value (padj) <0.05 and log(2) fold change >1 were considered as significantly differentially expressed. DGE was performed to compare the three clinical phenotypes: Late vs Early, Late vs Intermediate and Intermediate vs Early (Supplementary Figure S2c,d,e), and 1, 49 and 4 genes, respectively, were found to be differentially expressed in the three groups. It is likely that due to increased diversity of the signal in the extreme groups (i.e Early and Late), the small patient numbers and the stringency of the parameters, DGE was only able to return a few significant genes. When combining the patients into the larger groups, DGE identified four genes that were differentially expressed between non-SC versus SC (i.e rebound < vs > 500 days), and five genes between Progressors and PTCs (i.e rebound <vs > 100 days) (Supplementary Figure S2a,b). However, in view of these limitations we turned to alternative approaches.

We identified functionally linked ‘gene sets’ which can offer a more comprehensive insight than single genes into the mechanisms underpinning a phenotype. Gene Set Enrichment Analysis (GSEA) was performed on all genes ranked by DESeq2 Wald statistics (with the Wald test being applied to each gene), for each group of phenotypes. For the non-SC vs SC (i.e. rebound < vs > 500 days) analysis the gene set representing interferon signaling pathways was clearly enriched in the SC phenotype (Figure 1a). The PTC phenotype (rebound > 100 days) was associated with 51 gene sets and the most statistically significant enriched biological processes were related to immunoregulation, platelet activation and interferon alpha and beta signaling (Figure 1b). DGE and GSEA were also performed on the samples comparing the distinct phenotypes (Early, Intermediate and Late), and also showed interferon alpha and beta signaling to be enriched in the late rebounders (Supplementary Figure S3), however, as noted above, due to small sample sizes we were not able to take these analyses further.

**Figure 1:**
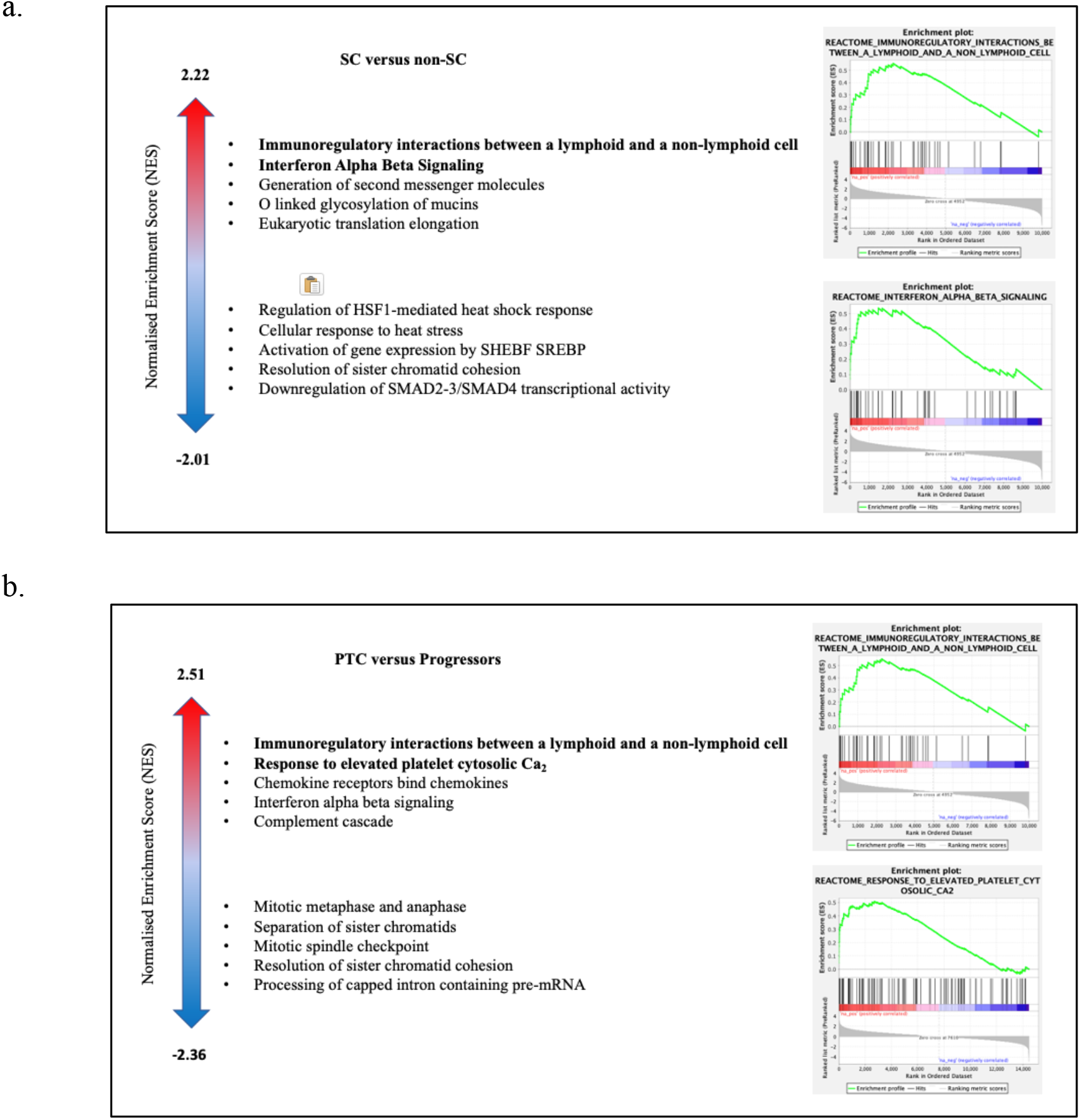
Pathway enrichment according to clinical phenotype using DESeq2 with GSEA and Rank Prod with ReactomePA.

To validate the results presented above, a non-parametric approach, RankProd, was employed as an alternative method for DGE followed by pathway enrichment with ReactomePA. This analysis also showed enrichment of type I Interferon pathways for the SC phenotype and for Platelet and Fc-gamma receptor (FCGR) activation pathways for PTCs compared to Progressors (Figure 1c-d), consistent with the GSEA.

### Weighted Gene Co-expression Network Analysis (WGCNA) identifies modules associated with rebound phenotypes and time to rebound

For an alternative interrogation of the transcriptomics data, Weighted Gene Co-expression Network Analysis (WGCNA) was employed to identify gene co-expression modules that relate to the clinical features of interest and inform on pathway enrichment. Aside from the different phenotype groups (SC versus non-SC and PTC versus Progressors), Time to Rebound was also used as a clinical trait of interest, as it offered the opportunity to explore the genetic correlations without having to make an a priori decision on sample grouping. WGCNA constructs a weighted network that represents the interaction patterns among genes, by emphasizing the strong gene-gene correlations at the expense of the weak ones in order to reduce noise. Here, the co-expression network was built using the expression data of a total of 4457 out of 28395 genes, that were retained after filtering for low variability. All pairwise correlations were calculated by raising the correlation values to the power of β=18, for which the scale-free topology fit index was 0.8 (Figure 2a). WGCNA then clustered all genes with similar co-expression patterns into modules, conventionally denoted by colour names (Figure S4). The modules were then associated with three external sample traits, namely SC (rebound >500 days), PTC (rebound >100 days) and ‘Days to Rebound’ as a continuous variable.

**Figure 2:**
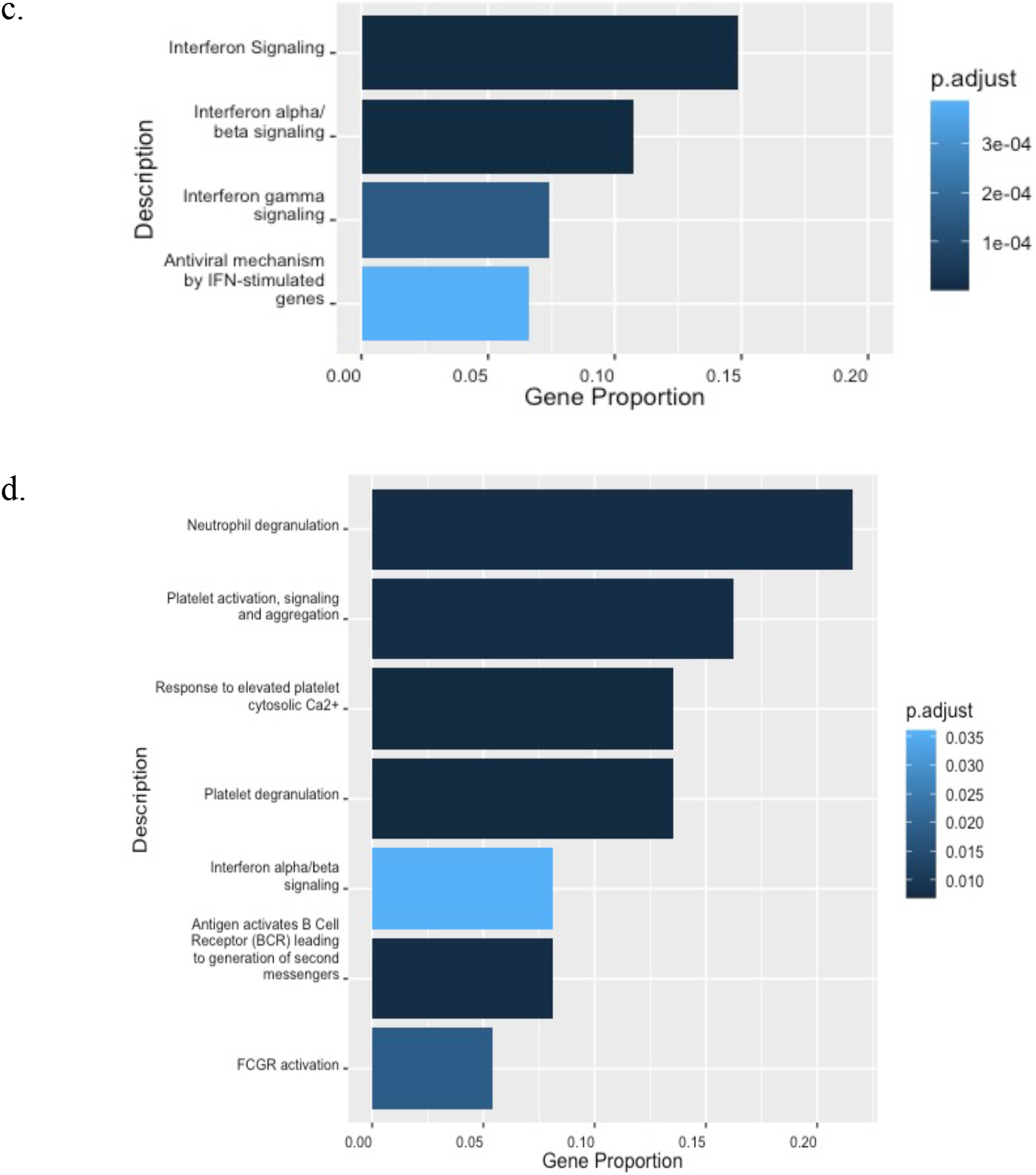
Pathway enrichment according to clinical phenotype using DESeq2 with GSEA and Rank Prod with ReactomePA. Top five up- and down-regulated pathways and GSEA pre-ranked enrichment plots for pathways (FDR<0.25) for SC versus non-SC phenotypes (a) and PTC versus Progressors (b). The range of Normalised Enrichment Scores (NES) shown on the left. Bar-plots presen plots presenting ReactomePA pathway enrichment for SC versus nonSC (c) and PTC versus Progressors (d).

Four modules were positively correlated with the traits of interest. ‘PaleTurquoise’ (labelled as Module 1 in Figure 2b) was correlated with both Time to Rebound (cor=0.64, p=0.02) and the SC phenotype (cor=0.62, p=0.03). ‘Salmon’ (Module 2; cor=0.58, p<0.05) and ‘Yellow’ (Module 3; cor=0.63, p=0.03) correlated with the SC phenotype. ‘Grey60’ (Module 4) was found to be associated with the PTC phenotype (cor=0.65, p=0.02) (Figure 2b). A significantly high correlation of Module Membership (MM) and Gene Significance (GS) is reported for these modules showing that the genes that are significantly associated with the respective traits are also important elements of each module (Figure 2c-g).

### Functional Enrichment of WGCNA modules associated with SC phenotype and Days to Rebound indicates Interferon Type I pathway involvement

The modules obtained from WGCNA were then uploaded on STRING for downstream enrichment analysis (Figure 3a-b). To be consistent with the GSEA analysis, the Reactome database was used to report the enrichment. The Module 1 genes (PaleTurquoise) showed an enrichment (p<0.05) in the Interferon Type I pathway (Figure 3c). Based on the GS and MM scores, 17 genes in Module 1 were identified as hub genes. Genes in modules 2 and 3 that correlated with the SC phenotype were shown to have non-significant enrichment values (0.06 and 0.3, respectively), indicating that these identified random proteins are not well connected. Module 4 (Grey60) that was associated with the PTC phenotype showed an enrichment (p<0.01) in platelet activation and degranulation (Figure 3d).

**Figure 3:**
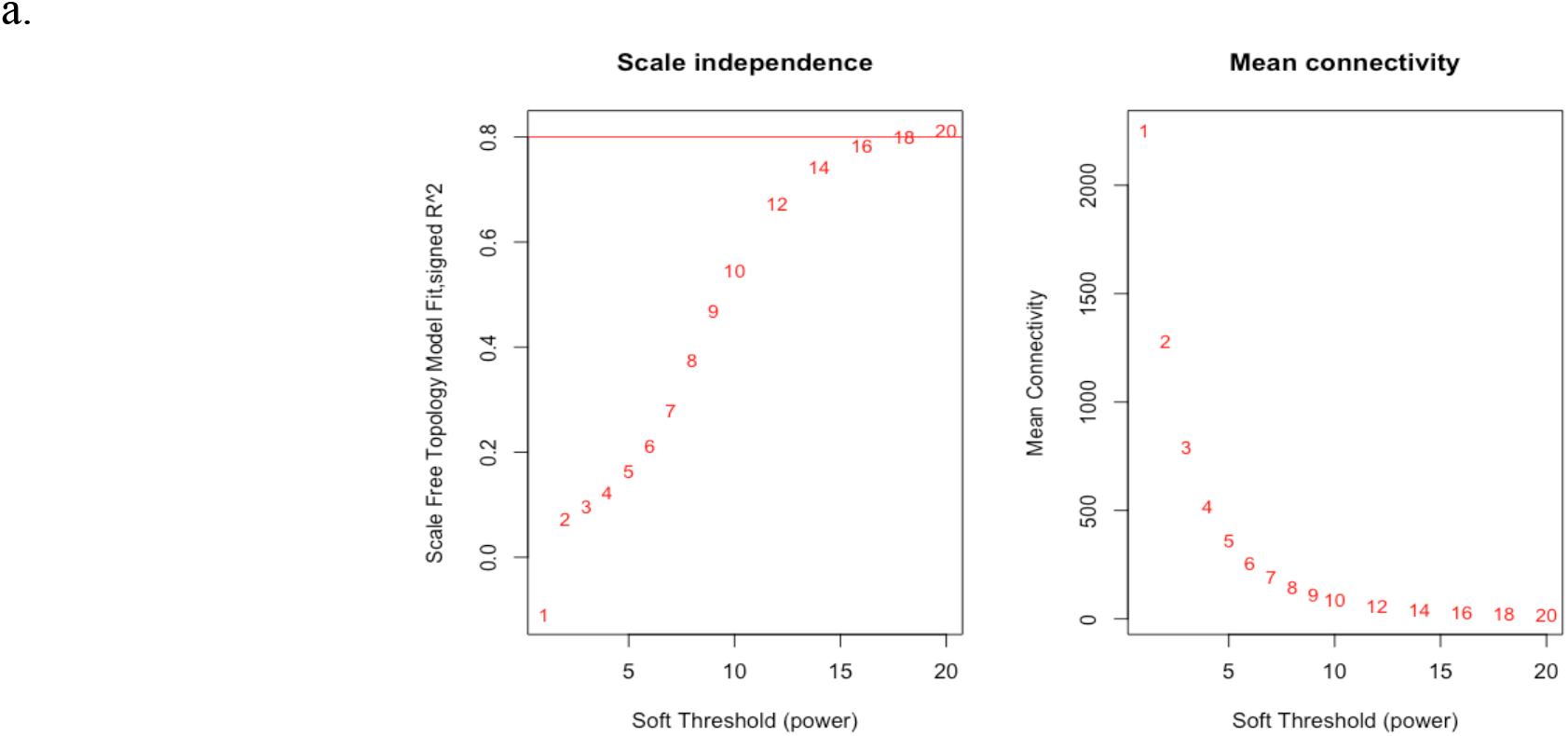
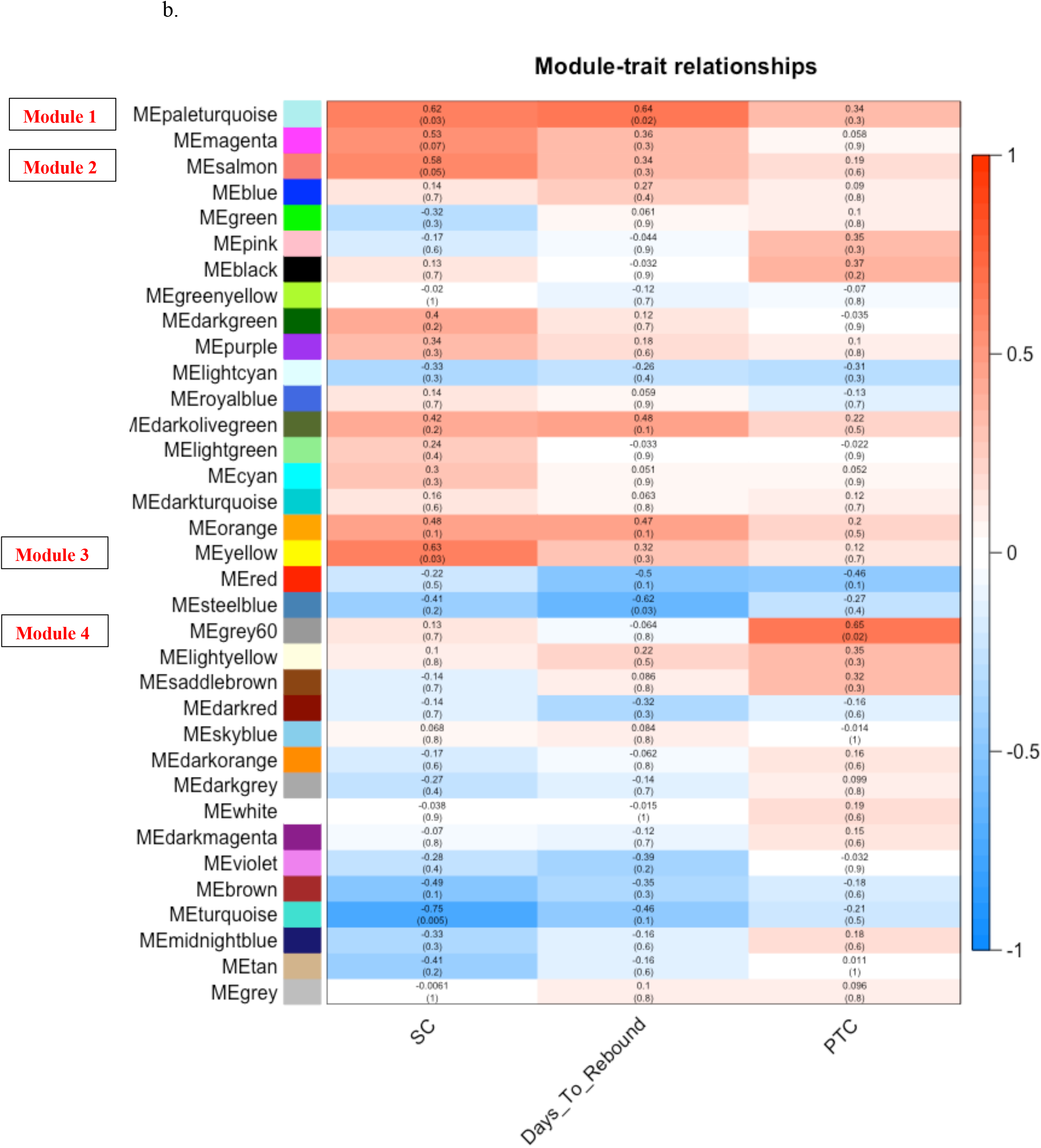
Network topology analysis and module identification using WGCNA.

### Risk score calculation based on the expression of two genes can potentially predict time to rebound

As a next step, we looked to see if any of the individual genes identified in the WGCNA results were more closely linked to progression, potentially leading to a signature that might predict longer post-TI viral suppression. Univariable analysis showed that the expression of 5 out of 17 hub genes contained in Module 1, namely ISG15, TRIM25, IFIT3, IFI6 and XAF1, was associated with a protective Hazard Ratio (HR) for early rebound <1 (Table 2).

**Table 2.** Univariable Cox Regression for Individual Protective Genes.

| Gene | HR | 95%CI | p-value |
| --- | --- | --- | --- |
| ISG15 | 0.11 | 0.02-0.66 | 0.01 |
| IFIT3 | 0.088 | 0.00-0.89 | 0.04 |
| TRIM25 | 0.021 | 0.00-0.75 | 0.03 |
| IFI6 | 0.074 | 0.00-0.84 | 0.02 |
| XAF1 | 0.19 | 0.04-0.90 | 0.03 |

HR= Hazard Ratio; 95% CI = 95% Confidence Intervals

Interestingly, all five are interferon stimulated genes. A multivariable Cox Regression analysis with LASSO was performed using the five significant genes and only TRIM25 and ISG15 were retained (Figure 4a). A Risk Score (RS) was calculated for all participants using the expression of these two genes and their LASSO coefficients (Table S2) were classified as either low or high relative to the mean RS. Kaplan Meier analysis showed that a lower RS with higher expression of both ISG15 and TRIM25 was found to be significantly associated with longer suppression post-TI, compared to participants with a high RS (P<0.001) (Figure 4b).

**Figure 4:**
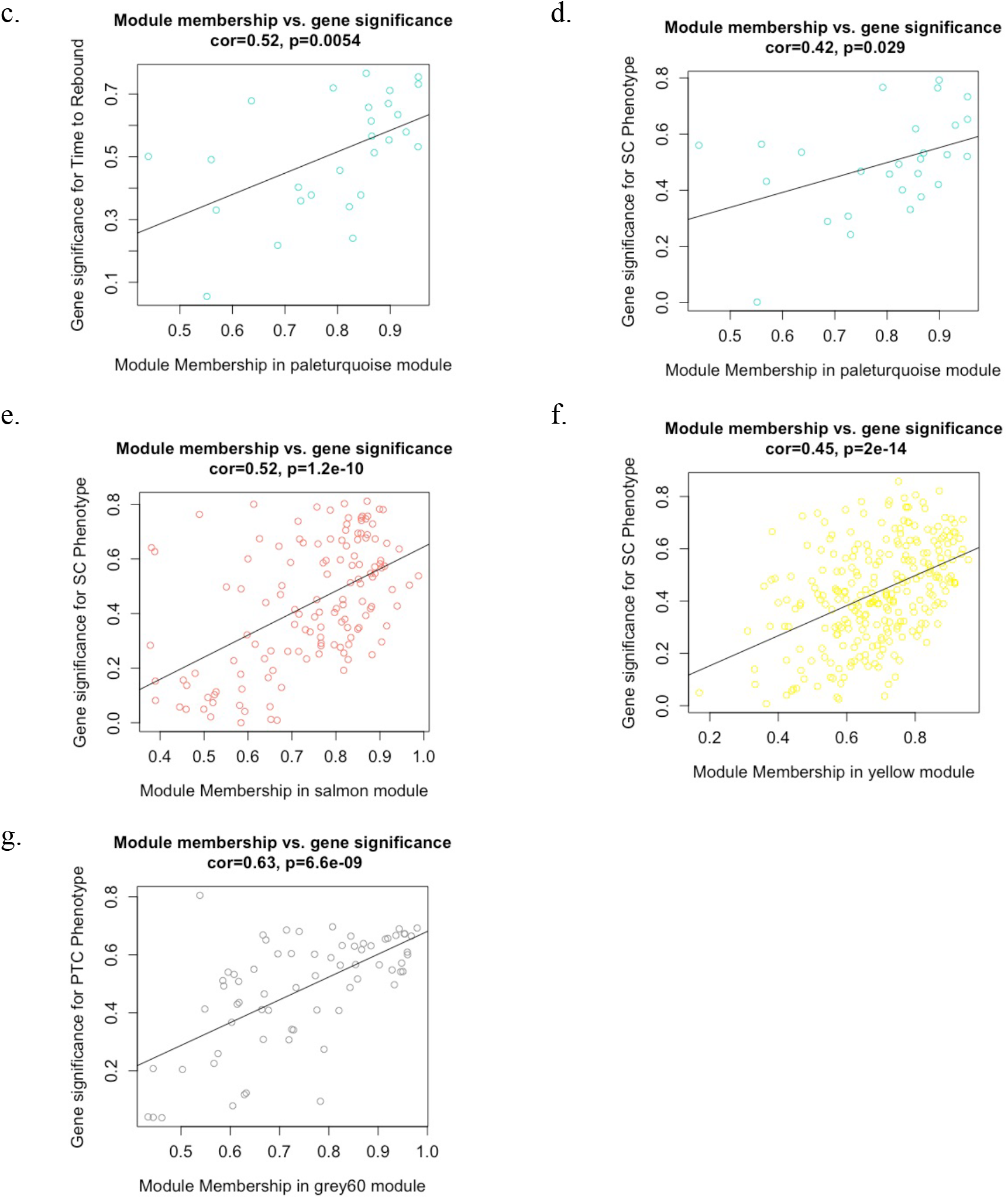
Network topology analysis and module identification using WGCNA. a. Network topology analysis for different soft thresholding powers. b. Heatmap plot of module-trait correlation and its statistical significance in parenthesis. The table is colour-coded based on direction and intensity of correlation, which is shown in red boxes. c-g: Scatterplots of gene significance (GS) for trait of interest versus module membership (MM): paleturquoise (days to rebound) (c); paleturquoise (SC phenotype) (d); salmon (SC phenotype) (e); yellow (SC phenotype) (f); grey60 (PTC phenotype) (g).

**Figure 5:** Gene Network and Pathway Enrichment Visualisation of selected WGCNA modules.

**Figure 6:**
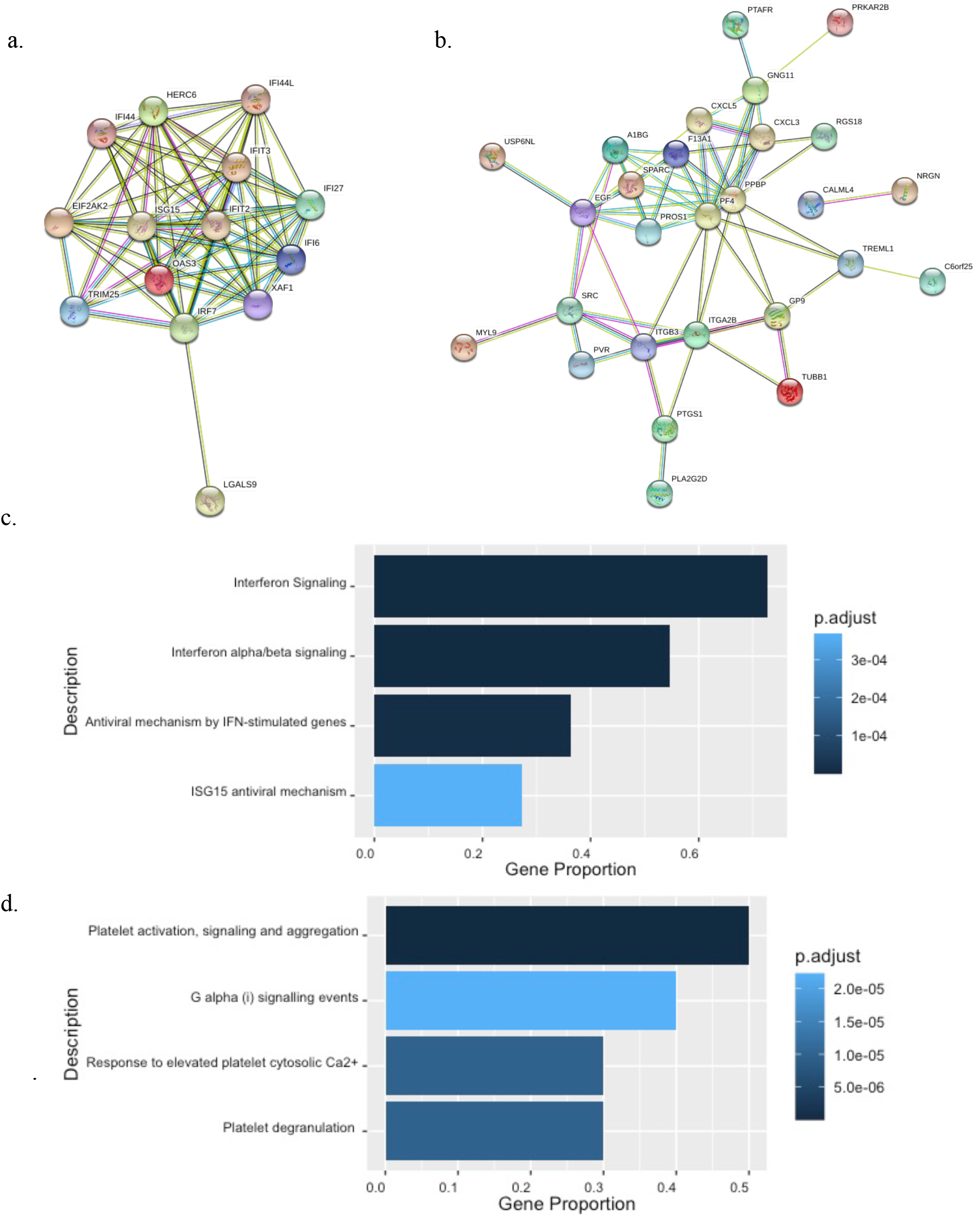
Gene Network and Pathway Enrichment Visualisation of selected WGCNA modules. Visualisation of the Protein-Protein Interaction network of genes within the PaleTurquoise (a) and the Grey60 (b) modules, generated by STRING. c-d. Top pathways enrichment bar plots for PaleTurquoise (c) and Grey60 (d) hub genes.

**Figure 7:** The expression of ISG15 and TRIM25 predict risk of earlier viral rebound.

**Figure 8:**
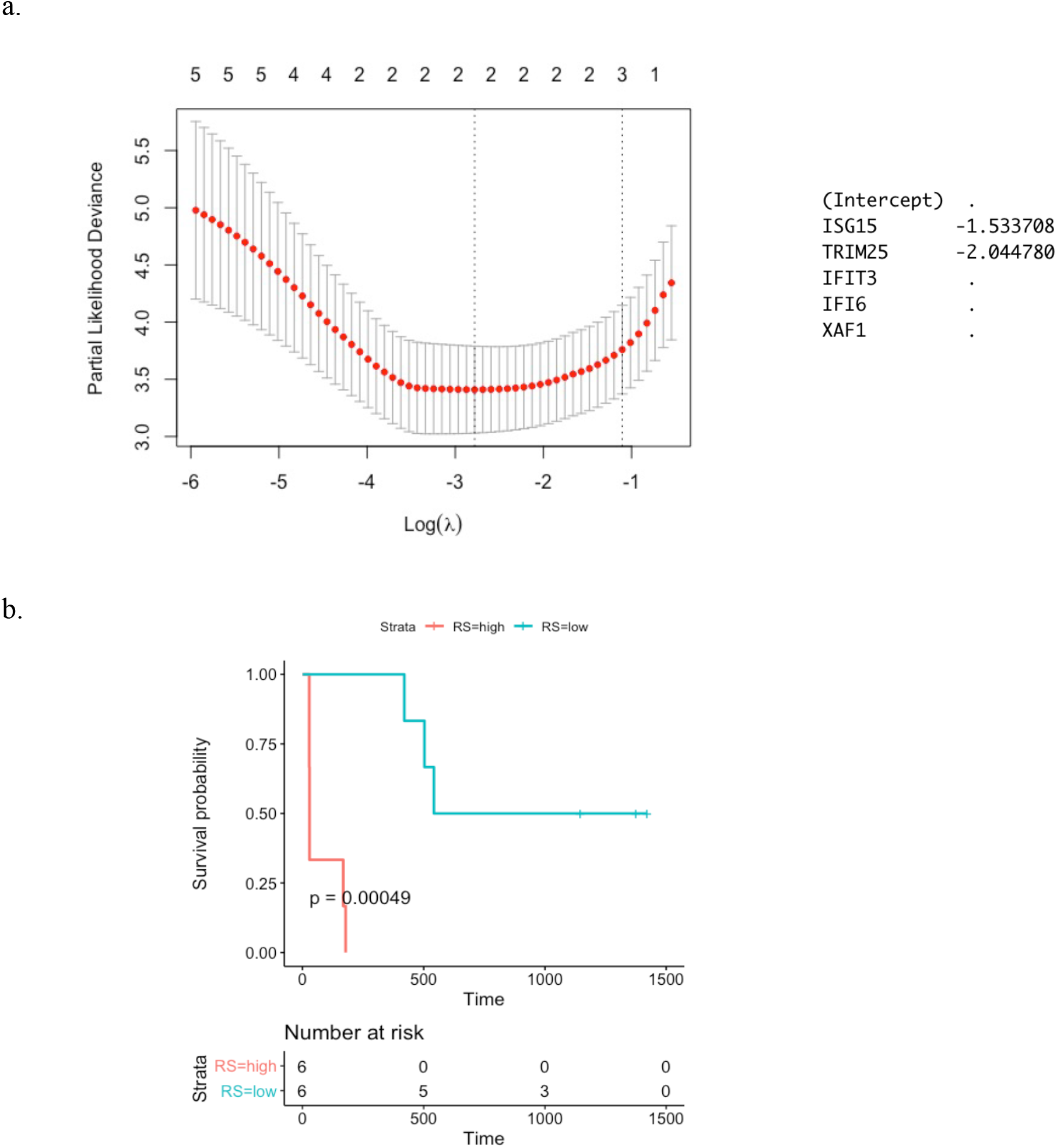
The expression of ISG15 and TRIM25 predict risk of earlier viral rebound. a. Estimation of λ and coefficient penalisation for gene selection with LASSO. Two of the five genes are selected as correlating with a lower risk of early rebound. Negative coefficient indicates the effect is protective. b. Kaplan-Meier curve of gene expression-based Risk Score (RS), predicting the likelihood of early and late post-TI rebound. Blue and red lines represent low and high-risk score respectively and ‘+’ marks represent the censored samples.

## Discussion

The majority of HIV positive individuals tend to rebound quickly after stopping ART, although it is evident that for a few some degree of virological remission is established. Understanding the mechanisms of post-treatment control through the identification of biomarkers that would help stratify patients into viral rebound risk groups would be invaluable for clinical decision making. Molecular studies constitute a powerful tool for identifying signatures that determine different response phenotypes. High-throughput RNA sequencing and subsequent bioinformatics analysis render an excellent opportunity to characterise the host immune profile of ART responders and provide a mechanism for identifying biomarkers that predict control post-ART cessation.

Our aim was to gain a deeper understanding of how host gene expression affected the duration of post-TI viral control using samples collected during the SPARTAC trial. This cohort offered a unique opportunity to study samples from participants with PHI that have been followed-up for an average of 4 years post-TI(Fidler et al., 2013). The samples used in this study were collected at the day of TI. To minimise the noise caused by averaging the expression of different cell types, only CD4+ T cells were selected and sequenced. This was because based CD4+ T cells comprise the majority of the HIV reservoir, and therefore any variability in transcriptional activity might impact the likelihood of proviral expression and rebound viraemia, and also related to the significance of the CD4+ T-cell response in the control of HIV(Frater et al., 2014). In addition, we restricted our analyses to women from Africa to exclude potential confounding factors from sex or ethnicity.

The results of DGE/GSEA revealed a distinct transcriptomic difference in those who controlled for over 500 days (i.e. between the SC phenotype and the non-SC phenotype), which was associated with the Type I Interferon Pathway (IFN-I). IFN-I pathways comprise a family of cytokines playing a dual role in the regulation of the innate immune response in infection(McNab et al., 2015). While continuous exposure to IFN-I inflicts chronic immune activation and subsequent exhaustion of T-cells and progression to AIDS, studies on SIV models illustrate the benefit of IFN-I administration at the early stage of infection in inhibiting viral replication(Deeks et al., 2017; Sandler et al., 2014; Van der Sluis et al., 2020). Similarly, HIV develops strategies to evade IFN-I and to inhibit the functionality of the proteins regulated by IFN-I (Fenton-May et al., 2013), suggestive of a fundamental role of this pathway in viraemic control. Additionally, the presence of less fit, adapted viruses - well described for T cell immune escape(Roberts et al., 2015; Zimbwa et al., 2007) - could also contribute to the difference in IFN-I response observed in this study(Adland et al., 2020; Cohn et al., 2018; Iyer et al., 2017). By defining post-treatment control (‘PTC’) participants as those that rebounded more than 100 days after TI(Etemad et al., 2019), we identified an enrichment of platelet activation pathways as well as IFN-I. The strong statistical association with platelet activation is intriguing and possibly consistent with reports that release of chemokines by activated platelets can function as an extra barrier at the early stages of HIV infection(Solomon Tsegaye et al., 2013). However, further investigation is required to determine the role of platelets in the defense against HIV.

WGCNA was then performed to determine how gene modules associate with specific clinical traits. For an unbiased analysis, ‘time to rebound’ was included with the SC and PTC phenotypes, to examine whether WGCNA findings corresponded with the DGE/GSEA analysis. The analysis identified two dominant modules of which one was associated with the ‘Time to Rebound’ and the SC trait, and one with the PTC phenotype. The gene ontology and pathway analysis showed a clear module enrichment in IFN-I associated with time to rebound, supporting the previous results for the phenotype traits. The majority of genes reported as hub genes were interferon stimulated genes (ISG), again consistent with the argument that the response to type I interferons is impacting rebound. A univariable Cox analysis identified five ISGs associated with remission in the participants (Table 2). Based on a multivariable Cox regression with LASSO screening of the five genes, a risk score based on the expression of ISG15 and TRIM25 only was developed to predict the likelihood of viral rebound. Higher expression of both genes was strongly protective for post-TI rebound. ISG15 has been reported to contribute to the inhibition of HIV virion release from infected cells(Gargan et al., 2018; Perng & Lenschow, 2018) and TRIM25 is an E3 ligase that positively regulates ISG15 conjugation to pathogen proteins, in a process with reported antiviral effects called ISGylation(Martín-Vicente et al., 2017). Although more work is underway to characterise the role of this pathway, this preliminary independent identification of these two genes would be consistent with a role in maintaining virological remission.

The small sample size on which this analysis was performed is a key limitation. In addition, our analysis was limited to bulk CD4+ T cells at a single timepoint, with no enrichment for those cells that were HIV-specific or contained proviral DNA. At this stage, the latter is not possible due to technological limitations. The other factor to consider is whether our choice of clinical phenotypes accurately discriminated between ‘post-treatment controllers’ and ‘elite controllers’. The latter have been well characterised(Martin et al., 2017), and more generally associated with effective HLA Class I-restricted T cell responses(Kiepiela et al., 2007), whereas it is still unclear as to which mechanisms are driving PTC. Larger studies will be needed to tease out these differences. However, that three independent analyses conferred the same statistically significant findings should be taken into consideration. Furthermore, this is a transcriptomic snapshot from the time of TI, and information about the expression of ISG is not available for other timepoints, before and after TI. This finding of a strong type I interferon signal associating with delayed rebound in this small study needs to be confirmed in larger clinical trials incorporating TI. If these data are reproducible, they could help with the identification of a valuable biomarkers of remission and pathways to drug discovery for the HIV cure field.

## Materials and Methods

### Participant characteristics and trial design

We analysed 18 SPARTAC participants who had received 48 weeks of suppressive ART commenced shortly after seroconversion. Participants were selected based on availability of viable peripheral blood mononuclear cells (PBMCs) and stratified by time to rebound. Participants were sampled at the time of TI, 48 weeks after commencing ART during Primary HIV Infection (PHI). The demographics of the participants are presented in Table 1, and Supplementary Table S1. Based on time to rebound, participants were classified as those who rebounded Early (<100 days after TI), Intermediate (100-500 days after TI) and Late (>500 days after TI). Initial analyses revealed gene expression varied significantly dependent on country of origin (African vs Non-African) (Supplementary Figure S1), and so - to avoid confounders - participants were analysed separately according to origin. Additionally, there were no male participants in the Late category, and so analyses including this parameter were restricted to female participants.

### RNA Isolation and Sequencing

CD4+ T cells were isolated using negative selection (Stem Cell Technologies CD4 enrichment kit) according to the manufacturer’s recommendations, and samples were processed in one batch. Total RNA was extracted using the Qiagen RNA Kit with Qiashredder columns and was sent for library preparation and sequencing. Material was quantified using RiboGreen (Invitrogen) on the FLUOstar OPTIMA plate reader (BMG Labtech) and the size profile and integrity analysed on the 2200 TapeStation (Agilent, RNA ScreenTape). Input material was normalised to 100ng prior to library preparation. Polyadenylated transcript enrichment and strand specific library preparation was completed using NEBNext Ultra II mRNA kit (NEB) following manufacturer’s instructions. Libraries were amplified on a Tetrad (Bio-Rad) using in-house unique dual indexing primers(Lamble et al., 2013). Individual libraries were normalised using Qubit, and the size profile was analysed on the 2200 TapeStation. Individual libraries were normalised and pooled together accordingly. The pooled library was diluted to ~10 nM for storage. The 10 nM library was denatured and further diluted prior to sequencing. Paired end sequencing was performed using an Illumina Novaseq6000 platform at 150 paired end mode (Illumina, San Diego, CA).

### Differential Gene Expression Screening and Gene Set Enrichment Analysis

DESeq2(Love et al., 2014) was used to compute the differential gene expression (DGE) between phenotypes using a featurecounts table. Only genes with a DGE of adjusted p-value of <0.05 were considered statistically significant. Gene Set Enrichment Analysis (GSEA)(Mootha et al., 2003; Subramanian et al., 2005) was used to detect differences in biologically relevant pathways in the dataset. The datasets were pre-ranked by the DESeq2 Wald statistic value. Permutation was set at 1000. Gene sets in the Reactome database(Jassal et al., 2020) were used as reference. Results satisfying a False Discovery Rate (FDR) cut off of <25% were considered statistically significant. Due to the small number of replicates, RankProd(Del Carratore et al., 2017) – an alternative nonparametric statistical method to identify differentially expressed genes - was also used. This method ranks the genes that consistently rank high as up- or downregulated in a number of experiments. The test was run on data transformed with the DESeq2 variance stabilising transformation. The percentage of false prediction (pfp) cut off was set to 0.25. The statistically significant upregulated genes for each phenotype, ranked by an increasing RP score, were further analysed with the ClusterProfiler(Yu et al., 2012) and ReactomePA(Yu & He, 2016) R packages, for pathway enrichment.

### Co-Expression Network Construction

Weighted Gene Co-expression Network Analysis (WGCNA)(Langfelder & Horvath, 2008; B. Zhang & Horvath, 2005) is an R package used for gene expression profiling and was applied to the identification of genes associated with time to rebound. After pre-processing for low variance filtering and outlier removal, an appropriate soft-threshold power was selected to promote and penalise the strong and weak gene connections, respectively. Following this, a signed network was created using the one-step approach, according to the package guidance. Genes were organised in modules, based on a common expression pattern. The expression of the genes in each module was summarised with an eigengene value, and a colour-label attributed to each to assist identification.

### Identification and Annotation of Important Modules and Hub genes

Module eigengenes were correlated with the clinical traits, in order to identify the most biologically relevant modules. The modules that were selected for downstream analysis were the ones that had a significant correlation with the trait. The most connected genes (hub genes), which are of functional significance, were defined by their Module Membership (MM>0.8) and their Gene-trait Significance (GS>0.2). All selected modules were uploaded on STRING(Szklarczyk et al., 2019) for pathway enrichment, gene ontology annotation and PPI visualisation.

### Identification of predictor genes for time to rebound

Survival analysis was performed for the identified hub genes, using the survival(Terry, 2020) package on R, to assess the prognostic value of the hub genes. The first viral rebound after TI was used as the event of interest. A univariable Cox Proportional Hazard Regression was performed on each gene. Genes with a statistically significant correlation to time to rebound were selected for a multivariable Cox regression with LASSO after selecting an optimal regularization parameter λ, to penalise the variable coefficients. A Risk Score (RS) to predict prognosis was calculated according to the formula below, where β is the multivariate Cox coefficient and exp is the expression value for all significant genes.

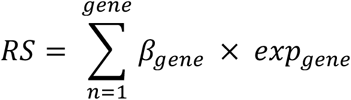

All participants that did not report post-TI rebound during the follow-up period of SPARTAC were classified as censored. All genes with a statistically significant association with a longer remission were then used to calculate a prediction score for time to rebound, by multiplying the gene coefficient with the gene expression.

## Acknowledgements

We thank the participants of SPARTAC and the SPARTAC Trial Investigators (listed below). The SPARTAC trial was funded by a grant from the Wellcome Trust (069598/Z/02/Z). JF is supported by the Medical Research Council (MR/L006588/1), and the National Institute for Health Research Oxford Biomedical Research Centre. PZ is supported through an NDM Studentship. This work was also supported by a British HIV Association (BHIVA) Research Award.

We acknowledge the role of Gita Ramjee (deceased) in the recruitment of participants to this study.

SPARTAC Trial Steering Committee: Independent Members-A Breckenridge (Chair), P Clayden, C Conlon, F Conradie, J Kaldor*, F Maggiolo, F Ssali, Country Principal Investigators-DA Cooper, P Kaleebu, G Ramjee, M Schechter, G Tambussi, JM Miro, J Weber. Trial Physician: S Fidler. Trial Statistician: A Babiker. Data and Safety Monitoring Committee (DSMC): T Peto (Chair), A McLaren (in memoriam), V Beral, G Chene, J Hakim. Co-ordinating Trial Centre: Medical Research Council Clinical Trials Unit, London (A Babiker, K Porter, M Thomason, F Ewings, M Gabriel, D Johnson, K Thompson, A Cursley*, K Donegan*, E Fossey*, P Kelleher*, K Lee*, B Murphy*, D Nock*). Central Immunology Laboratories and Repositories: The Peter Medawar Building for Pathogen Research, University of Oxford, UK (R Phillips, J Frater, L Ohm Laursen*, N Robinson, P Goulder, H Brown). Central Virology Laboratories and Repositories: Jefferiss Trust Laboratories, Imperial College, London, UK (M McClure, D Bonsall*, O Erlwein*, A Helander*, S Kaye, M Robinson, L Cook*, G Adcock*, P Ahmed*). Clinical Endpoint Review Committee: N Paton, S Fidler. Investigators and Staff at Participating Sites: Australia: St Vincents Hospital, Sydney (A Kelleher), Northside Clinic, Melbourne (R Moore), East Sydney Doctors, Sydney (R McFarlane), Prahran Market Clinic, Melbourne (N Roth), Taylor Square Private Clinic, Sydney (R Finlayson), The Centre Clinic, Melbourne (B Kiem Tee), Sexual Health Centre, Melbourne (T Read), AIDS Medical Unit, Brisbane (M Kelly), Burwood Rd Practice, Sydney (N Doong), Holdsworth House Medical Practice, Sydney (M Bloch), Aids Research Initiative, Sydney (C Workman). Coordinating Centre in Australia: Kirby Institute University of New South Wales, Sydney (P Grey, DA Cooper, A Kelleher, M Law). Brazil: Projeto Praca Onze, Hospital Escola Sao Francisco de Assis, Universidade federal do Rio de Janeiro, Rio de Janeiro (M Schechter, P Gama, M Mercon*, M Barbosa de Souza, C Beppu Yoshida, JR Grangeiro da Silva, A Sampaio Amaral, D Fernandes de Aguiar, M de Fatima Melo, R Quaresma Garrido). Italy: Ospedale San Raffaele, Milan (G Tambussi, S Nozza, M Pogliaghi, S Chiappetta, L Della Torre, E Gasparotto), Ospedale Lazzaro Spallanzani, Roma (G DOffizi, C Vlassi, A Corpolongo). South Africa: Cape Town: Desmond Tutu HIV-1 Centre, Institute of Infectious Diseases, Cape Town (R Wood, J Pitt, C Orrell, F Cilliers, R Croxford, K Middelkoop, LG Bekker, C Heiberg, J Aploon, N Killa, E Fielder, T Buhler). Johannesburg: The Wits Reproductive Health and HIV-1 Institute, University of Witswatersrand, Hillbrow Health Precinct, Johannesburg (H Rees, F Venter, T Palanee), Contract Laboratory Services, Johannesburg Hospital, Johannesburg (W Stevens, C Ingram, M Majam, M Papathanasopoulos). Kwazulu-Natal: HIV-1 Prevention Unit, Medical Research Council, Durban (G Ramjee, S Gappoo, J Moodley, A Premrajh, L Zako). Uganda: Medical Research Council/Uganda Virus Research Institute, Entebbe (H Grosskurth, A Kamali, P Kaleebu, U Bahemuka, J Mugisha*, HF Njaj*). Spain: Hospital Clinic-IDIBAPS, University of Barcelona, Barcelona (JM Miro, M Lopez-Dieguez*, C Manzardo, JA Arnaiz, T Pumarola, M Plana, M Tuset, MC Ligero, MT Garca, T Gallart, JM Gatell). UK and Ireland: Royal Sussex County Hospital, Brighton (M Fisher, K Hobbs, N Perry, D Pao, D Maitland, L Heald), St James’s Hospital, Dublin (F Mulcahy, G Courtney, S O’Dea, D Reidy), Regional Infectious Diseases Unit, Western General Hospital and Genitourinary Dept, Royal Infirmary of Edinburgh, Edinburgh (C Leen, G Scott, L Ellis, S Morris, P Simmonds), Chelsea and Westminster Hospital, London (B Gazzard, D Hawkins, C Higgs), Homerton Hospital, London (J Anderson, S Mguni), Mortimer Market Centre, London (I Williams, N De Esteban, P Pellegrino, A Arenas-Pinto, D Cornforth*, J Turner*), North Middlesex Hospital (J Ainsworth, A Waters), Royal Free Hospital, London (M Johnson, S Kinloch, A Carroll, P Byrne, Z Cuthbertson), Barts & the London NHS Trust, London (C Orkin, J Hand, C De Souza), St Marys Hospital, London (J Weber, S Fidler, E Hamlyn, E Thomson*, J Fox*, K Legg, S Mullaney*, A Winston, S Wilson, P Ambrose), Birmingham Heartlands Hospital, Birmingham (S Taylor, G Gilleran). Imperial College Trial Secretariat: S Keeling, A Becker. Imperial College DSMC Secretariat: C Boocock.

* Left the study team before the trial ended.

## Supplementary Materials

**Table S1:**
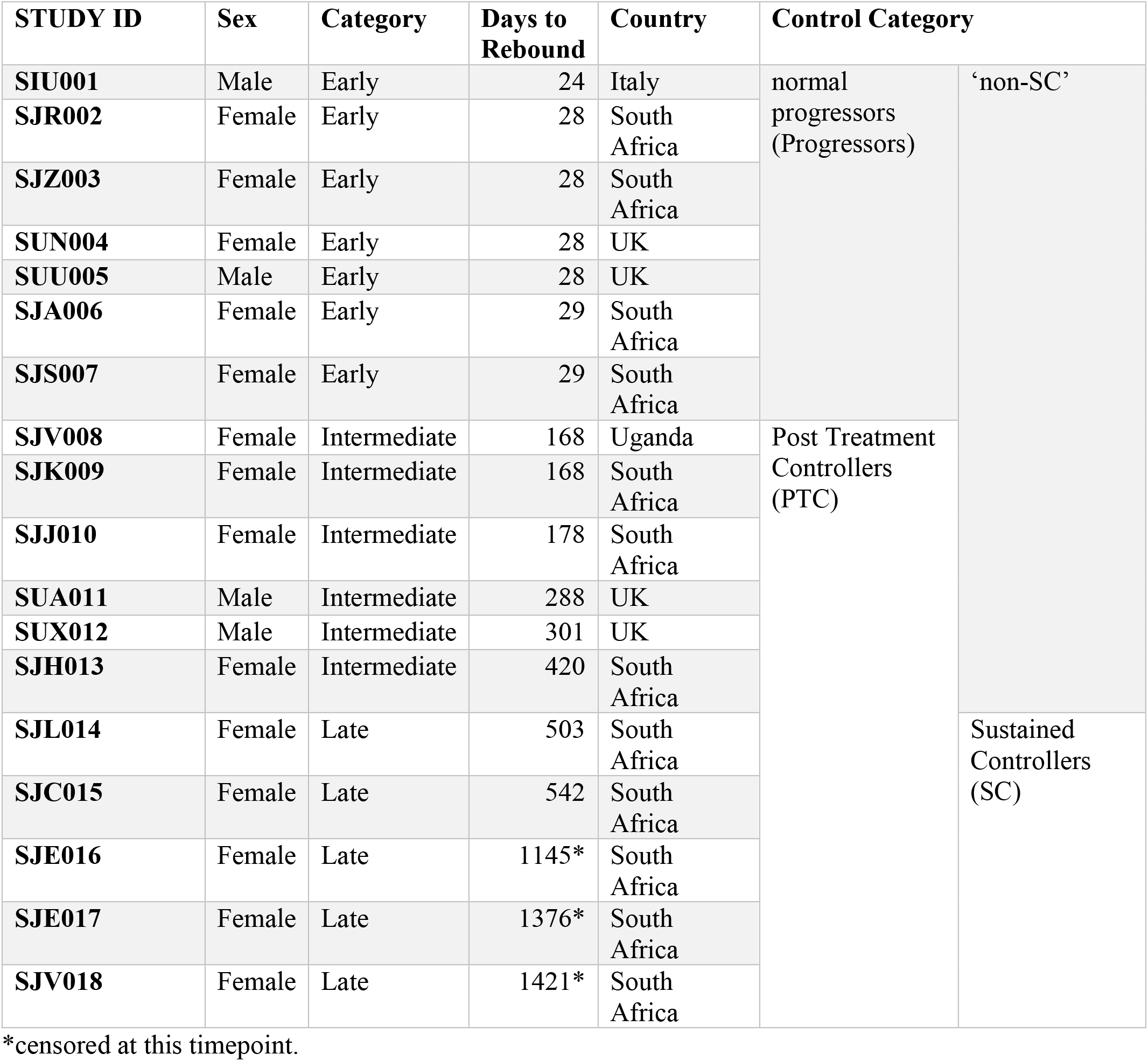
Detailed Demographics.

**Table S2:**
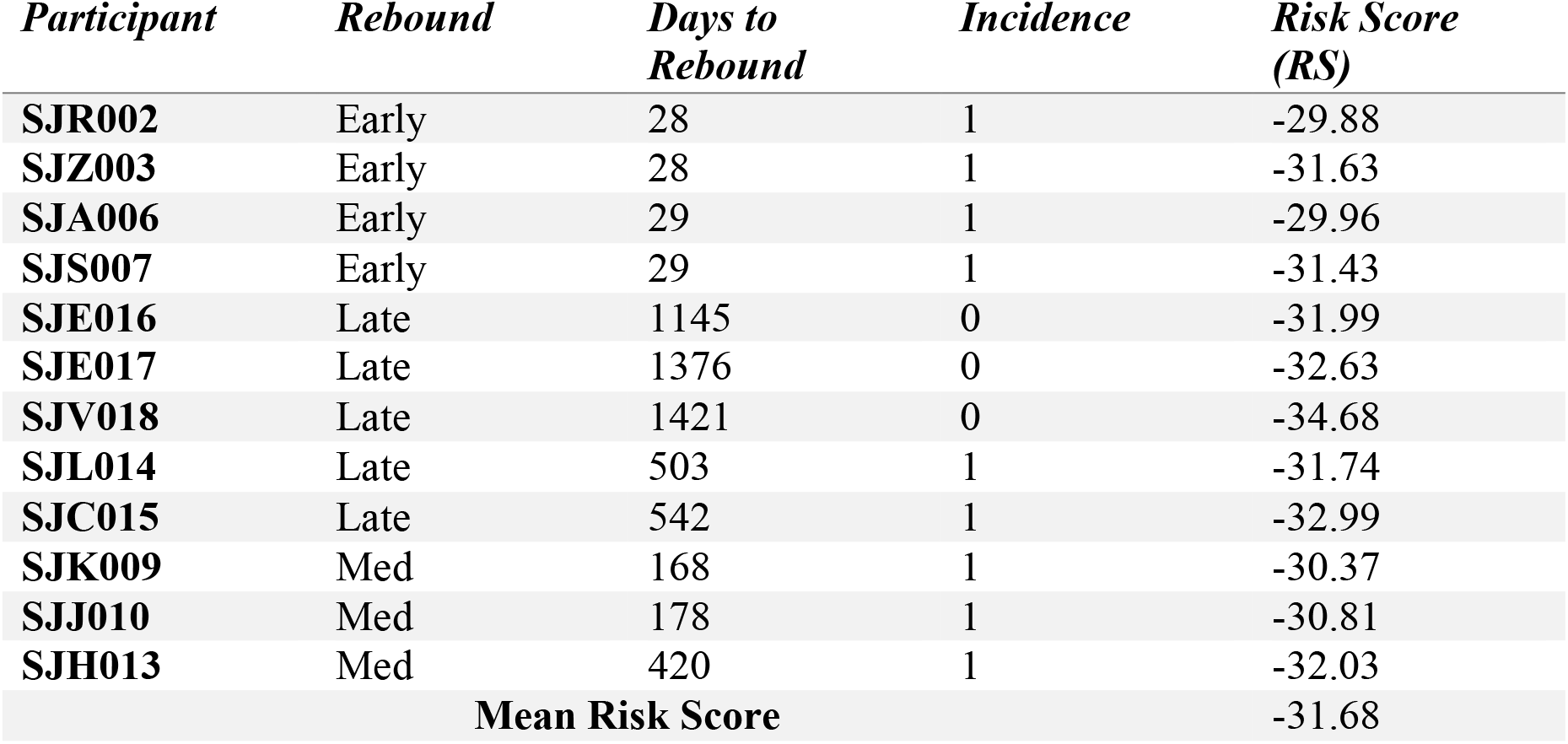
Individual Risk Score per participant, based on the expression of ISG15 and TRIM25

**Figure S1:**
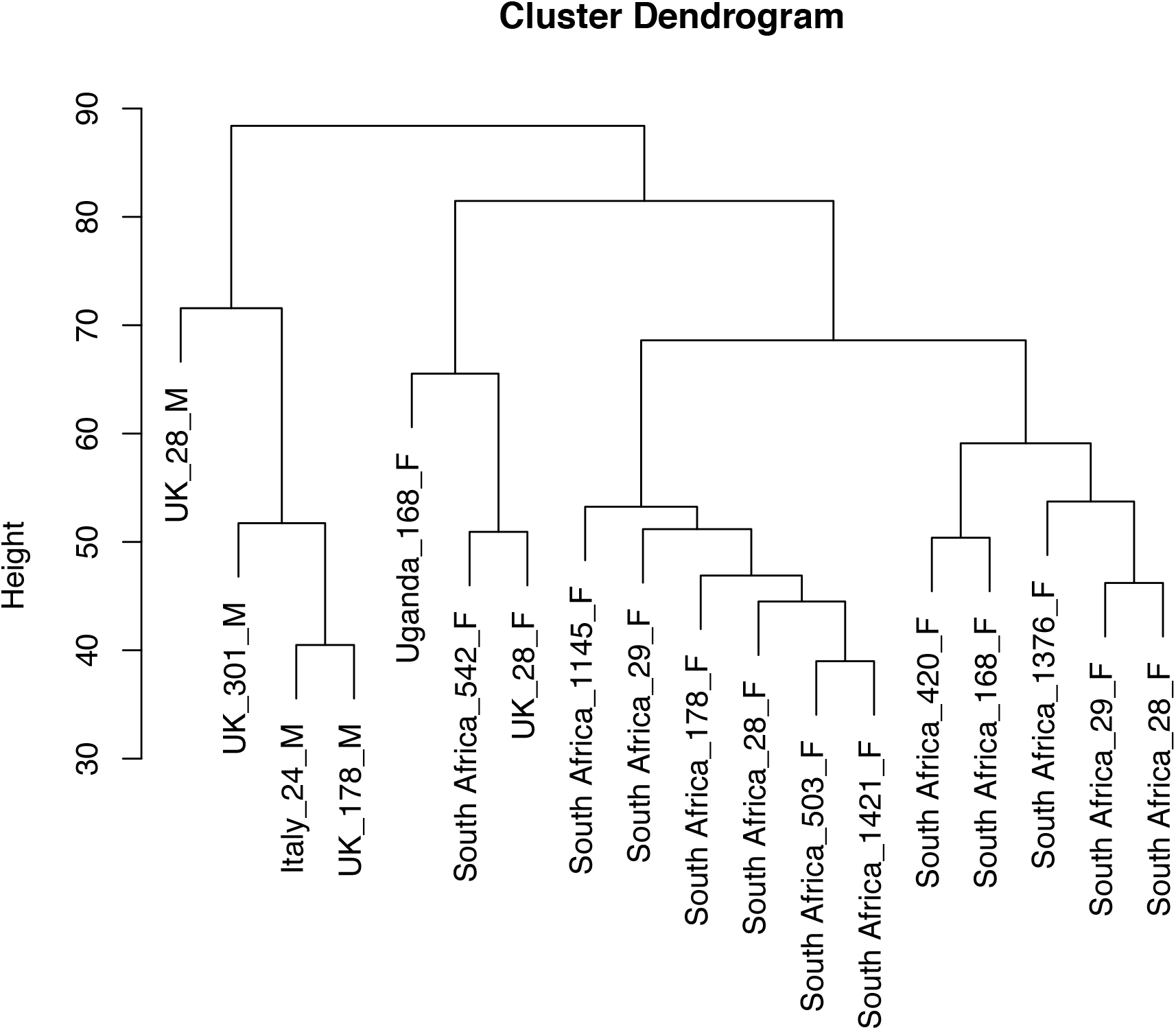
Dendrogram showing clustering of participants from different countries based on gene expression.

**Figure S2:**
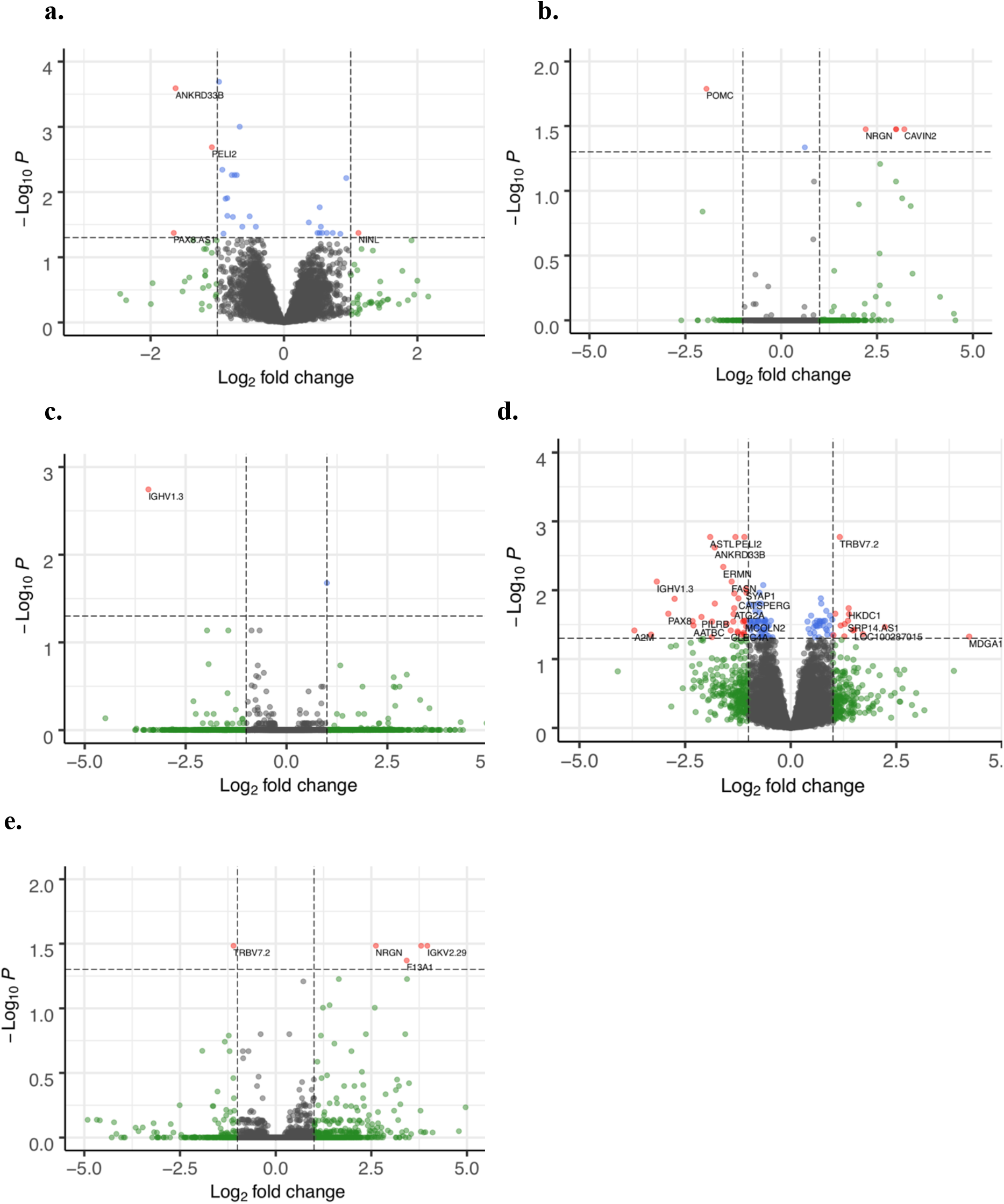
Volcano plots showing the differentially expressed genes according to patient clinical phenotype. SC versus nonSC (a), PTC versus Progressors (b), Late versus Early (c), Late versus Intermediate (d) and Intermediate versus Early (e). Red is used for genes significantly differentially expressed.

**Figure S3:**
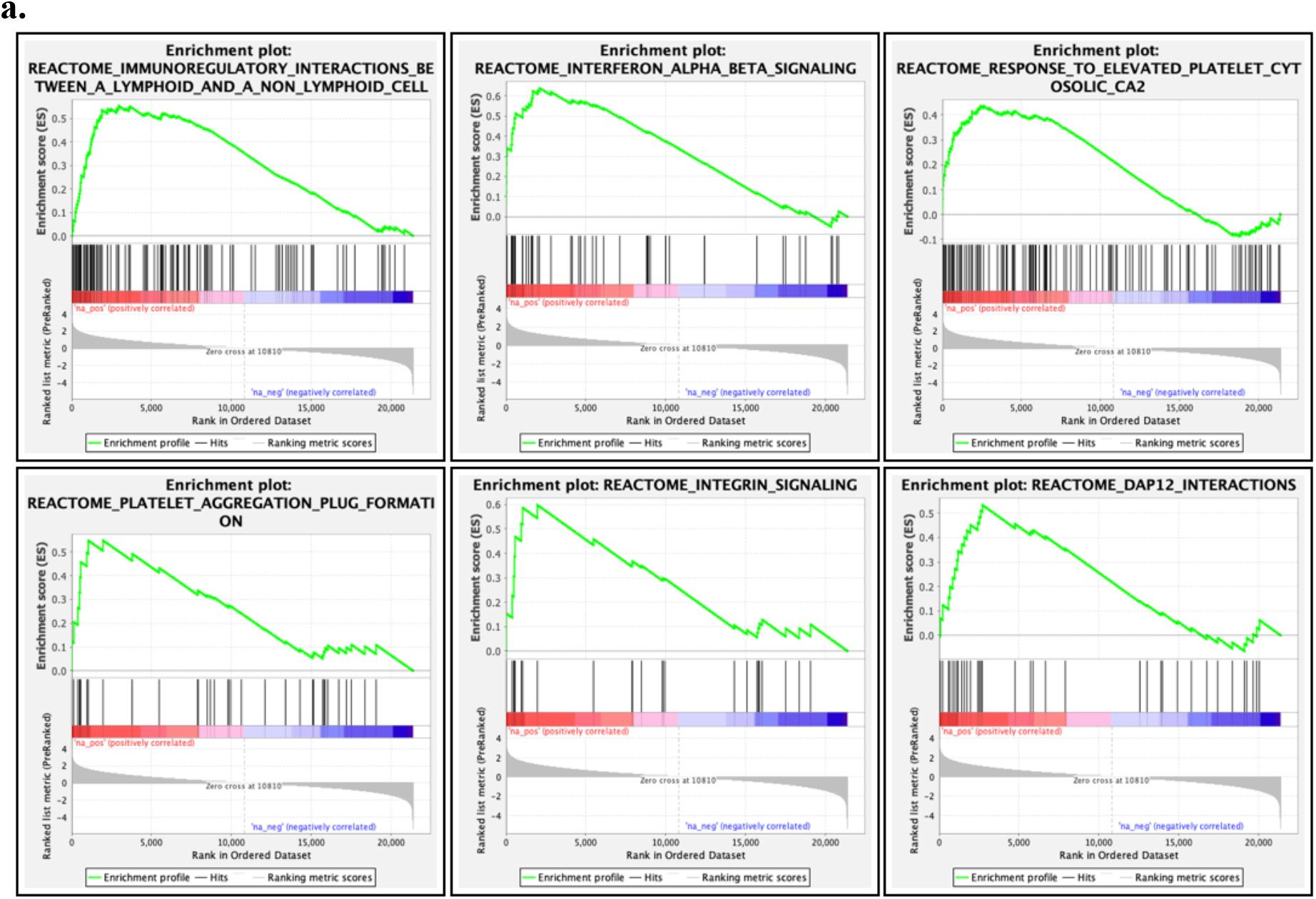
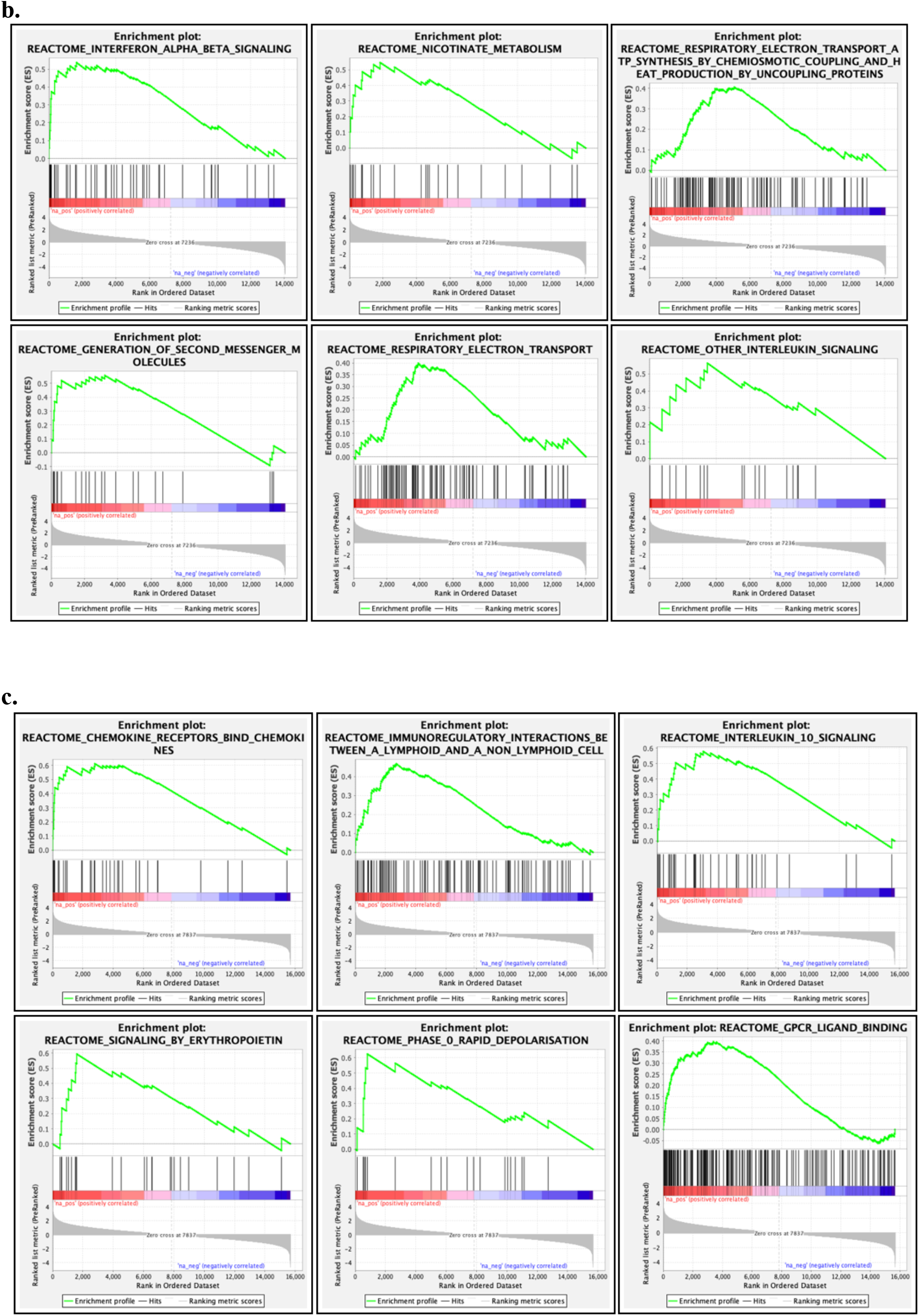
GSEA analyses according to clinical phenotype. top 6 pathways for Late versus Early (a), Late versus Intermediate (b) and Intermediate versus Early(c).

**Figure S4:**
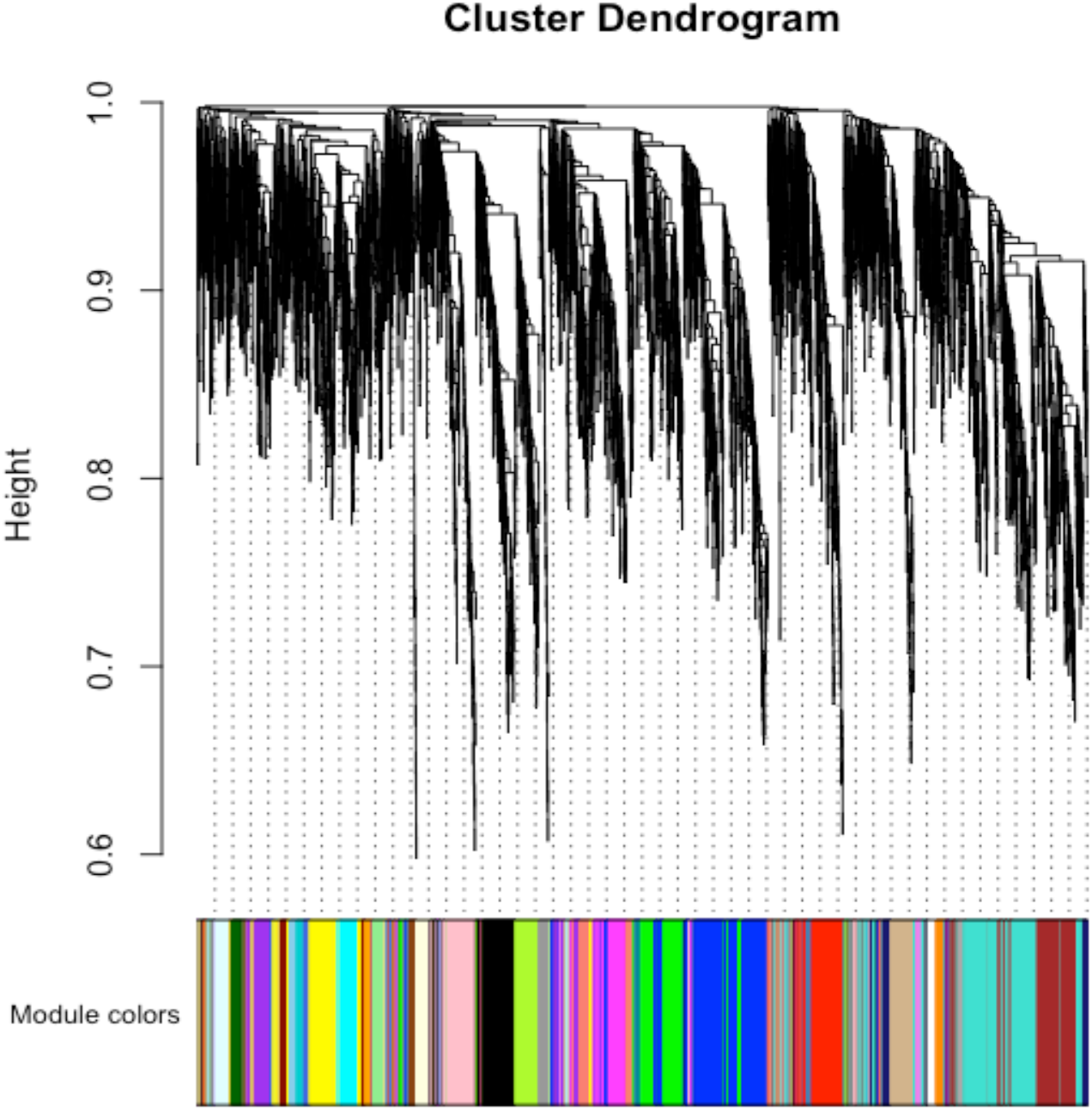
GSEA analyses according to clinical phenotype. top 6 pathways for Late versus Early (a), Late versus Intermediate (b) and Intermediate versus Early(c).

